# Before and after delisting: population dynamics of North Atlantic humpback whales over two decades in the Gulf of Maine

**DOI:** 10.1101/2024.02.04.577870

**Authors:** Jooke Robbins, Martine Bérubé, Phillip J. Clapham, David K. Mattila, Per J. Palsbøll, Regina Asmutis-Silvia, Alex Hill, Laura J. Howes, Scott Landry, Shelley Lonergan, Dianna Schulte, Jennifer E. Tackaberry, Mason T. Weinrich, Richard M. Pace

## Abstract

Whales are long-lived, slow-reproducing species that were decimated by commercial whaling. Although some populations seem close to recovery, others are challenging to assess. We studied humpback whales (*Megaptera novaeangliae*) in the Gulf of Maine, an area of the North Atlantic (NA) with well-documented threats. Long-term studies and mark-recapture data were used to estimate humpback whale apparent survival, abundance and population growth from 2000 through 2019. Estimates were derived from a hierarchical, Bayesian state-space model with sex, age, and random time effects on survival while accounting for individual capture probability. Abundance increased from 744 (95% CI: 726-762) in 2000 to 1,706 (95% CI: 1,639-1,771) in 2019, with 4.6% mean annual growth. However, adult males exhibited higher survival and outnumbered adult females by the end of the study. Over time, fewer calves were observed, calf survival varied and the juvenile class declined. These are rare insights into the dynamics underlying whale abundance trends and they revealed similarities to an endangered species that is declining due to environmental and human impacts. The results inform a listing change under the U.S. Endangered Species Act, a mortality event of unprecedented magnitude off the U.S and humpback whale recovery from historical whaling in the NA.

## 1. Introduction

Historical commercial whaling decimated whale species worldwide, including the humpback whale (*Megaptera novaeangliae*, Clapham et al. 1999; Rocha et al. 2015). Whereas some humpback whale populations are now thought to be at or close to recovery (Noad et al. 2019; Zerbini et al. 2019), the status of others is uncertain (Punt et al. 2006; Smith & Pike 2009; Thomas et al. 2016). In the U.S., the humpback whale was listed as an endangered species across its global range when the Endangered Species Act (ESA) became law in 1973. Over four decades later, the species was removed from the Endangered Species List and re-assessed for protection as 14 Distinct Population Segments (DPSs, NOAA 2016). The West Indies DPS, the primary breeding population of humpback whales in the western North Atlantic, is one of nine DPSs that no longer receives protection under the ESA (NOAA 2016). While the DPS is unlikely to be in jeopardy of extinction, there are nevertheless questions about its status of recovery from whaling and its current population dynamics. Notably, available abundance data have been insufficient to determine population trends since early 1990s (Bettridge et al. 2015; Hayes et al. 2018).

Humpback whales in the West Indies DPS breeding population maintain maternally directed fidelity to specific mid- to high-latitude feeding grounds (Martin et al. 1984; Clapham & Mayo 1987). The Gulf of Maine (GoM), along the northeastern coast of the United States and southeastern Canada, is the southwestern-most primary humpback whale feeding ground in the North Atlantic (Katona & Beard 1990; Stevick et al. 2006). The GoM is also an area with high human activity and the humpback whales that feed there are vulnerable to a range of unintentional anthropogenic impacts that have exceeded management limits (van der Hoop et al. 2013; Hayes et al. 2022). There have been four Unusual Mortality Events (UMEs) involving humpback whales in the GoM since the 1980s, including one event that has been ongoing since the DPS was removed from the Endangered Species List (Geraci et al. 1989; NOAA 2024). Humpback whale abundance and survival trends are essential to place these events in context, including in relation to population management and the mitigation of human impacts.

Long-term studies of individual humpback whales in the GoM have amassed life history and demographic data, the extent and detail of which are uncommon in other parts of the world. When integrated within a mark-recapture framework, such data can provide novel insights into the population processes that underlie abundance trends. Here, we used a combination of mark-recapture survey data, auxiliary sightings and demographic information in a Bayesian state-space mark-recapture statistical model to estimate humpback whale survival and abundance in the GoM over two decades, from 2000 through 2019. We assess an ESA recovery benchmark for the first time and provide insight into the effects of a UME that has been underway along the U.S. East Coast since the delisting. We also compare our findings to the dynamics of a declining and critically endangered species, the North Atlantic right whale (*Eubalaena glacialis*). Finally, we place our results in the context of population recovery from historical commercial whaling.

## 2. Materials and methods

Humpback whales can be individually identified based on their natural markings, particularly the shape and ventral pigmentation pattern of the flukes (Katona & Whitehead 1981). We obtained mark-recapture data from dedicated vessel surveys in the GoM (Figure 1), auxiliary sightings from an opportunistic sighting network and demographic data from longitudinal studies, as described in Appendix A. We estimated abundance using a mark-recapture statistical modeling approach similar to one employed to assess the population abundance trends of North Atlantic right whales, *Eubalaena glacialis* (Pace et al. 2017; Pace 2021). We used re-sighting histories of marked individuals to estimate survival rates and abundance in a Bayesian, state-space formulation estimated using Markov Chain Monte Carlo (MCMC) simulation. Specifically, we modified the approaches of Kéry & Schaub (2011) and Royle & Dorazio (2012) to produce a multi-state formulation that relied on Jolly-Seber open population model concepts of estimating the probability of new member entry (Jolly 1965; Seber 1982) but executed it in a Bayesian framework together with data augmentation. We separated the predictive relationships associated with state transition (or biological process) from those of the observation process and used auxiliary sightings to inform the biological state of individuals when they were missed by the survey. We then used logistic relationships with linear combinations of predictors to estimate apparent survival and capture probabilities for each age and sex class while accounting for sources of heterogeneity (Lebreton et al. 1992). When combined with an estimate of probability of entry, this allowed for the estimation of annual abundance (N_t_). Population growth rate was derived from the Bayesian state-space model outputs and the crude birth rate was calculated as the number of calves per individual each year as recorded in the primary dedicated surveys. Details of the Bayesian state-space model structure, implementation, derived products and statistical tests are provided in Appendix A.

**Figure 1.**
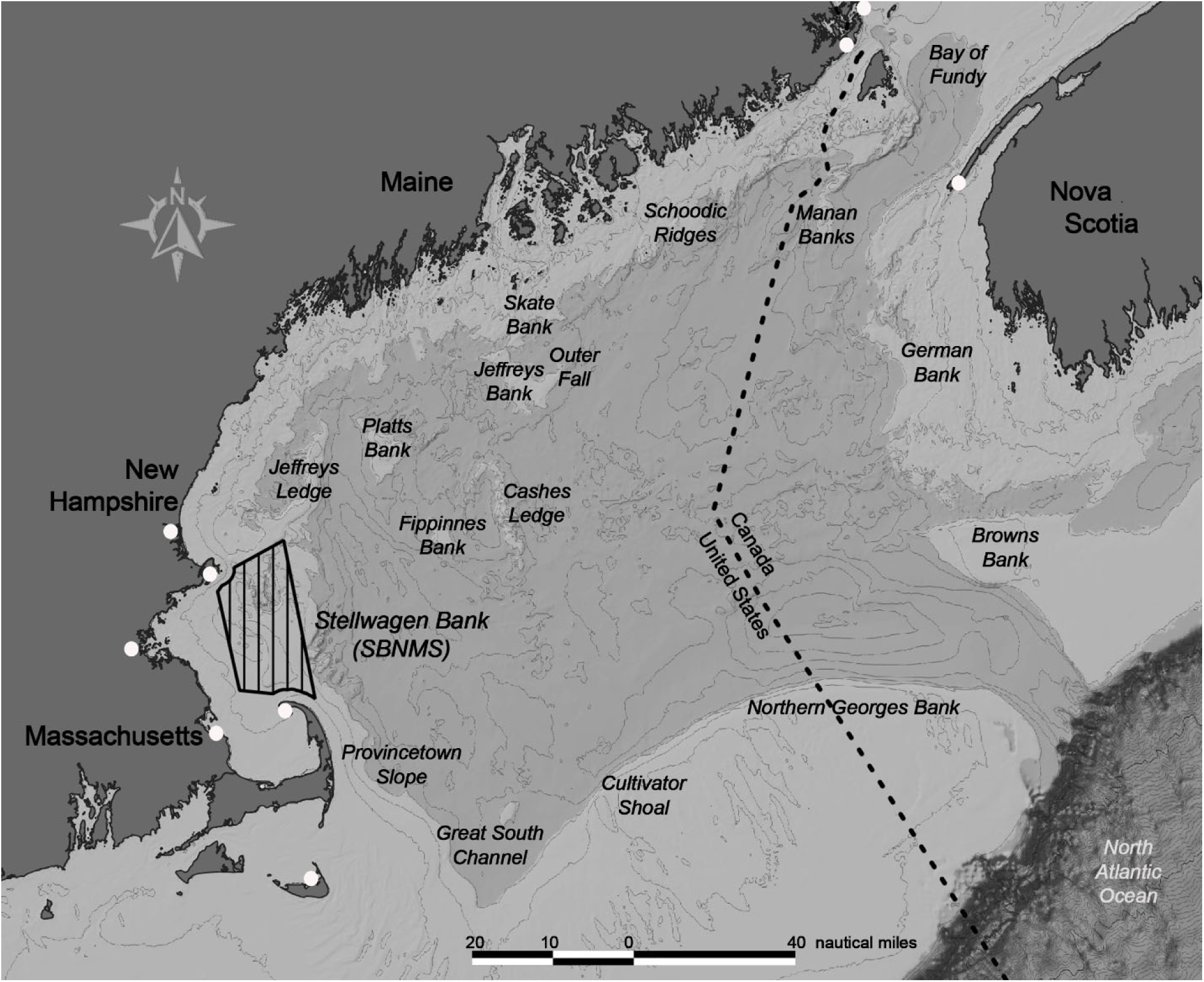
Gulf of Maine study area. Primary mark-recapture surveys focused on humpback whale aggregation sites (labeled in italics) around the inner margin of the GoM and Bay of Fundy, including Stellwagen Bank National Marine Sanctuary (hatched area). The white circles indicate the home ports of nine auxiliary data platforms.

## 3. Results

Between 2000 and 2019, the primary mark-recapture surveys documented 1,757 unique individuals and 6,401 individual sighting occasions over the years. An average of 320 individuals were seen per year (SD=96.9, Table B.1), and the average individual was seen in 3.6 years. Many individuals were seen only once (44.3%, n=778) and a minority (9.7%, n=171) were seen in more than half of the survey years (10-18 years). Most of the sightings occurred in the southwestern or northeastern GoM where auxiliary platforms (primarily commercial whale watching) were also clustered, but there was exchange among all of the areas where whales were found (Figure 2).

**Figure 2.**
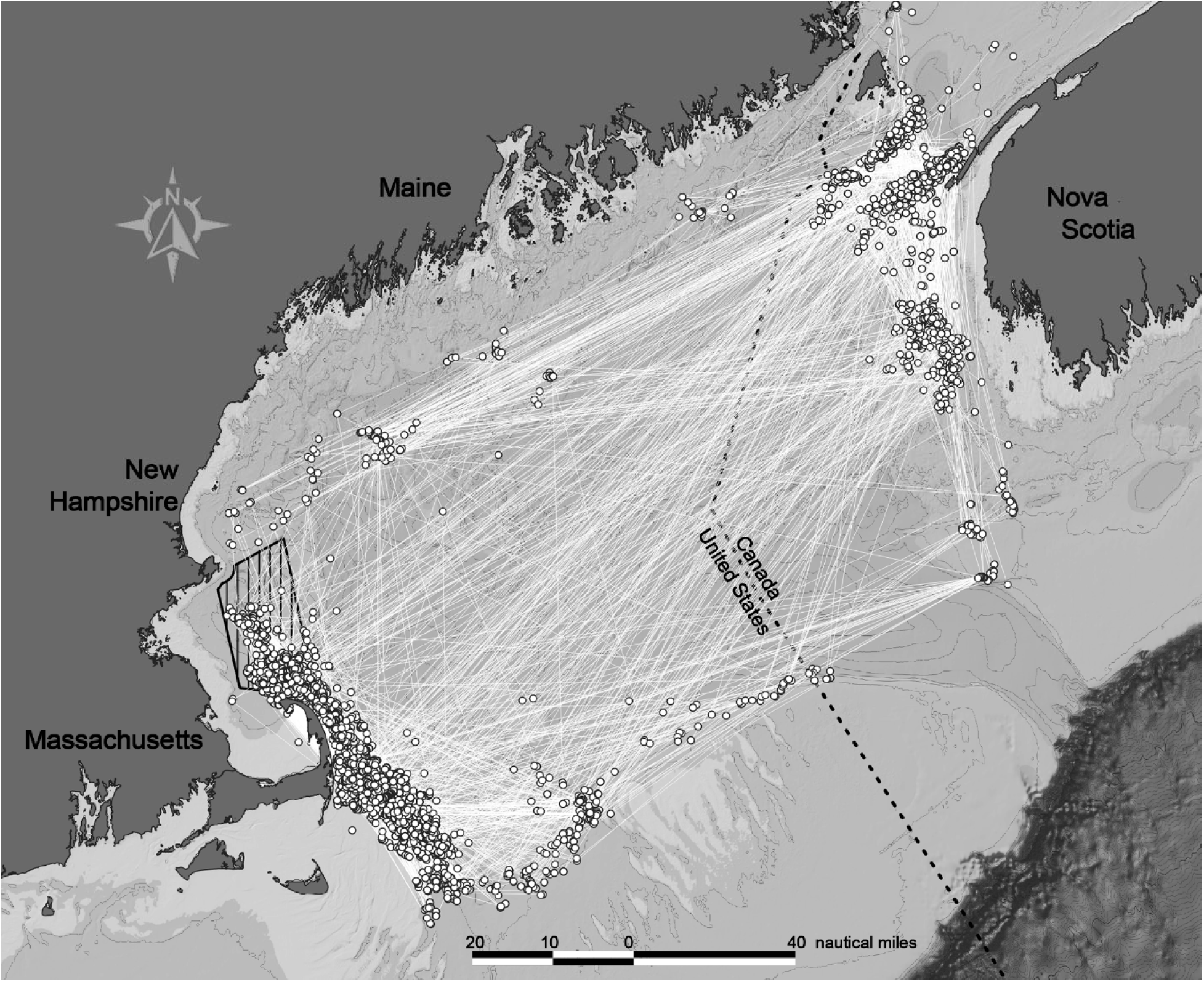
Primary mark-recapture sighting locations (white circles). The white lines connect the first sighting of an individual in a given year to its first sighting the next year (inter-annual displacement). There was individual exchange among all sampled areas.

The primary state-space Bayesian mark-recapture model had excellent Gelman-Rubin convergence statistics results and posterior distributions for all linear (logistic) parameters associated with time. Sex and age covariates contributed significantly to estimates of capture probability and survival (i.e., they were distinct from zero). Capture probabilities achieved by the primary mark-recapture surveys varied by year and were relatively low in most years except 2003 when we performed the survey twice in the same season (Figure B.1). The inclusion of auxiliary data on biological state increased the estimated abundance in all years, but especially in the last few intervals (Figure B.2).

Total abundance estimated from the state-space model increased from 744 (95% CI: 726-762) individuals in 2000 to 1,706 (95% CI: 1639-1,771) in 2019. (Figure 3). The population was estimated to number 1,589 (95%CI: 1,531-1,644) in 2016 when the West Indies DPS was removed from the U.S. Endangered Species List. Partitioning abundance by age class suggested that adults of both sexes increased in number across the study period. However, adult male survival rates were higher than females (Figure 4) and they numbered 13% (n=121) more than females by the end of the study (Figure 3). Juvenile survival was lower than for adults in all years and especially for calves in the year after weaning (Figure 4). Juvenile survival and abundance varied over time but the portion of the population known to be less than 5 years old declined by 60% between 2000 and 2019. The crude birth rate varied over time (mean=0.104, SD=0.036) but declined significantly over this period (p=0.0054, tau= -0.308, 95%CI: -0.6348, - 0.3189, Figure 5). The average rate of population growth was estimated to be 4.6% per year, but it was negative or flat during three periods of low apparent survival (2005-2006, 2011-2013, 2016) and was anomalously high (15.8%) in 2002 (Figure 6).

**Figure 3:**
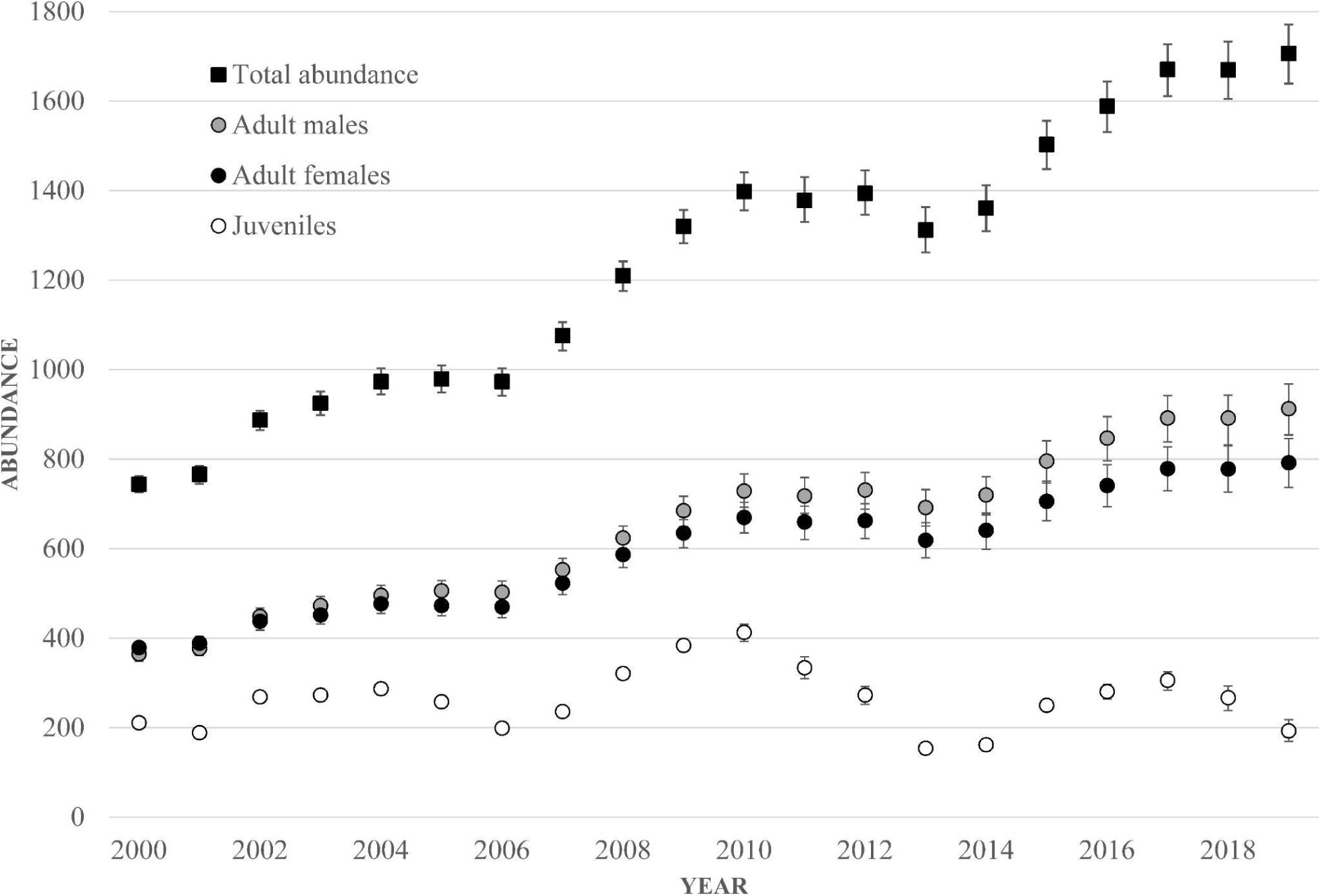
Abundance estimates and their 95% credible intervals for all whales (black squares), adult females (black circles), adult males (gray circles) and juveniles (white circles) in the Gulf of Maine from 2000 through 2019.

**Figure 4:**
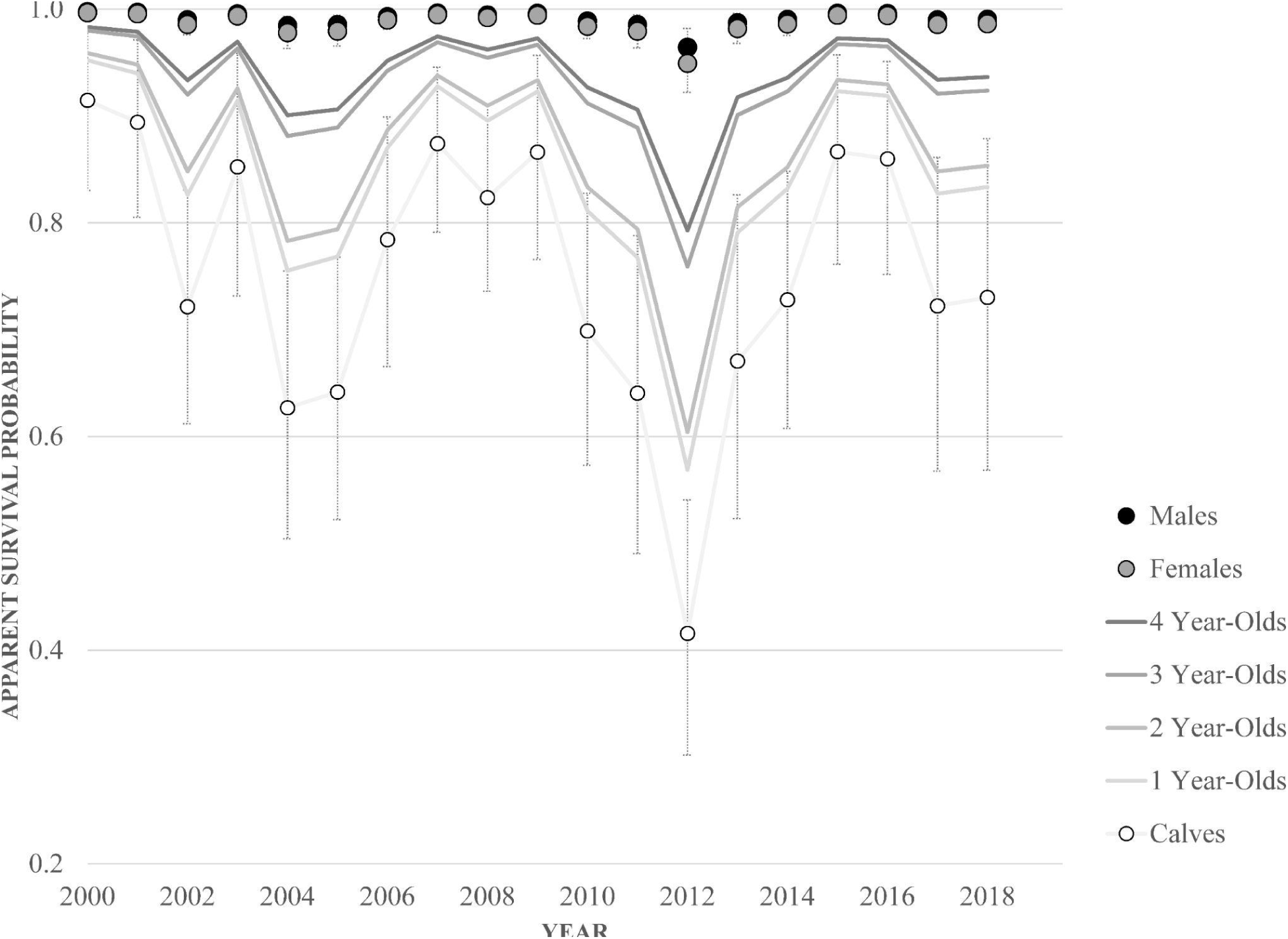
Apparent survival estimates and their 95% credible intervals for adult males (black circles), adult females (gray circles), and calves (<1 year old, white circles) from 2000 through 2019. Apparent survival of older juveniles (ages 1-4) are shown as lines only.

**Figure 5:**
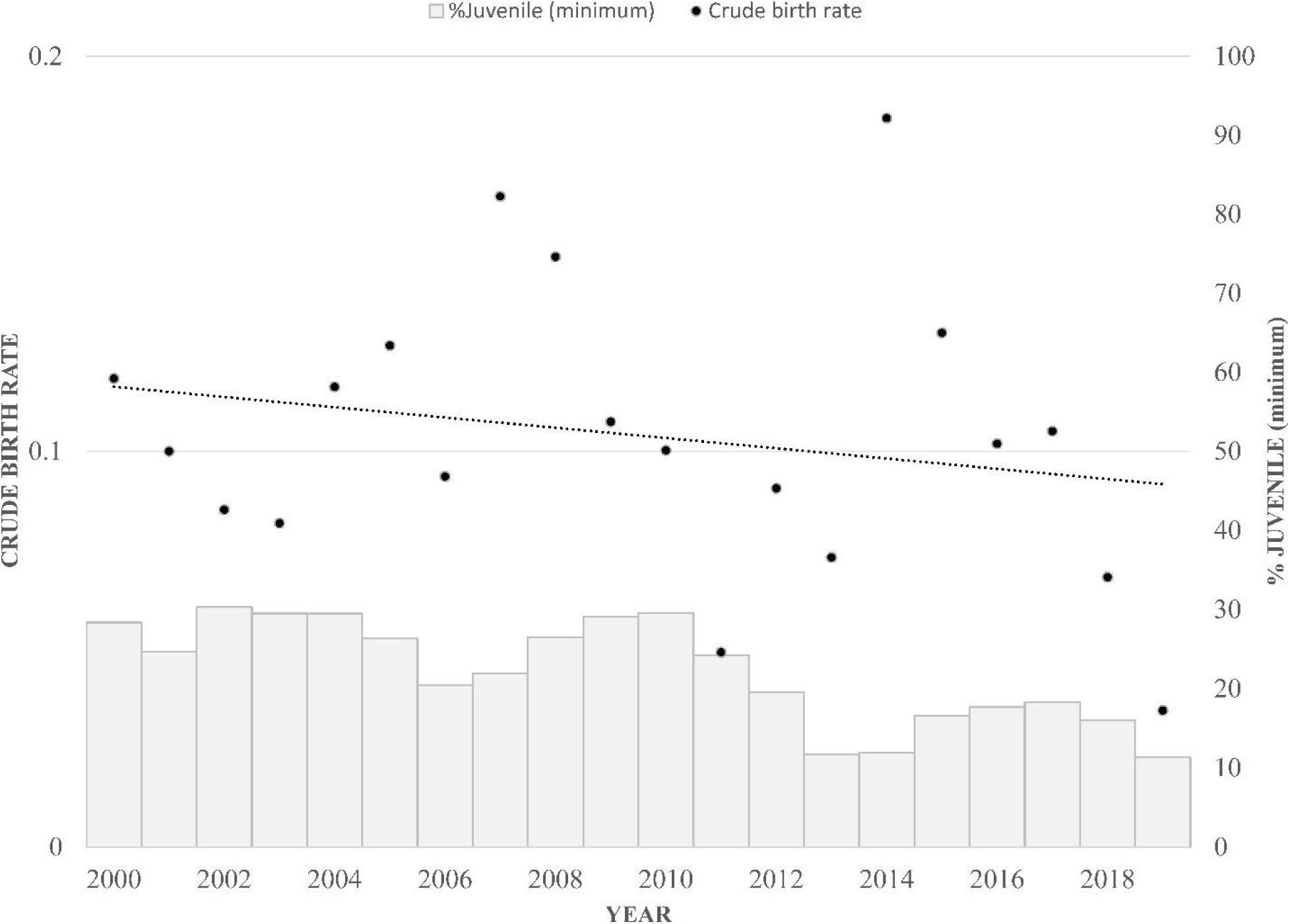
Crude birth rate (black circles with linear trend line) and minimum percentage of juveniles in the population (gray bars, right axis).

**Figure 6:**
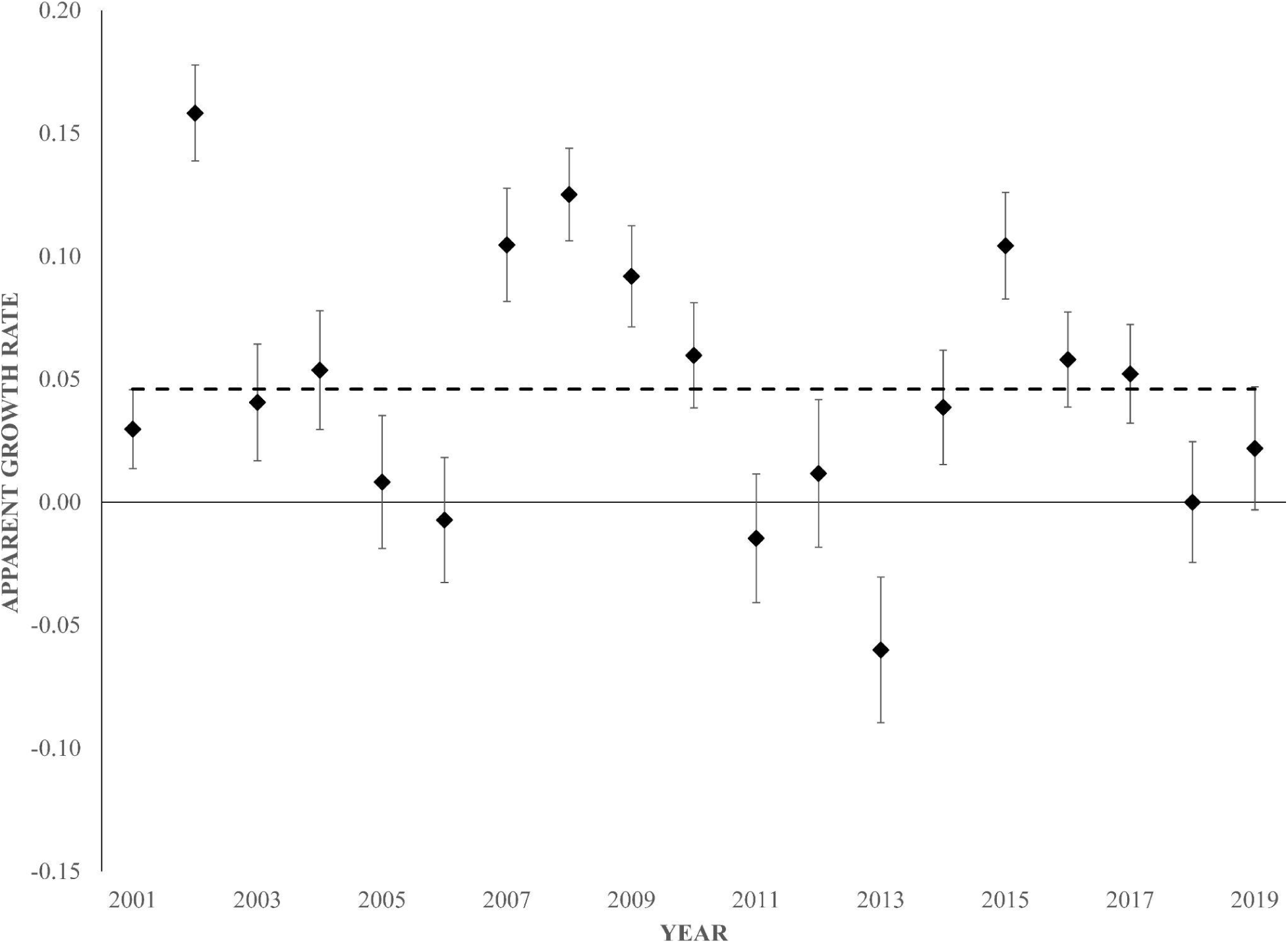
Apparent humpback whale population growth in the Gulf of Maine, 2001-2019, as derived from the Bayesian state-space model. Median growth rate with standard deviation for pairs of adjacent years. The dashed line represents the mean growth rate (4.6%).

## 4. Discussion

Information on population abundance and trends is essential for understanding population dynamics and managing accordingly. However, whale populations are challenging to monitor over appropriate spatial scales and timeframes due to their long lifespans, high mobility, slow reproductive rates, and their heterogeneous spatial distribution. They were decimated by commercial whaling and remain vulnerable to unintended impacts from even the most common human activities at sea, such as vessel operation and fishing. Here, we estimated humpback whale abundance and trend for the southwestern-most primary feeding ground in the North Atlantic using a combination of directed mark-recapture research, an auxiliary data-collection network and demographic data from more than 40 years of longitudinal research. We employed a novel mark-recapture statistical approach to maximize the use of these data, as well as account for heterogeneity and uncertainty. The results are the first extended time series of abundance in decades for any humpback whale feeding ground in the western North Atlantic.

### 4.1 Population dynamics of humpback whales in the Gulf of Maine

The results of this study indicate that the overall number of humpback whales using the GoM has increased over the past two decades. However, adult female abundance appears to have lagged behind that of adult males in recent years, and the minimum number of juveniles varied over time with no net increase between 2000 and 2019. This is the first study to partition whale abundance by age class and the decline in juvenile abundance is consistent with the observed low and variable calf survival rates, as well as the decline in calving rates over the study period.

The age- and sex-stratified apparent survival patterns underlying these estimates were consistent with the results of previous studies on this population (Robbins 2007), although the methods used here differed somewhat from prior Cormack-Jolly-Seber type analyses. Age-specific survival has not been estimated for other humpback whale populations, but our results are consistent with age-specific patterns of survival for long-lived mammals more generally (Caughley 1966, Gaillard et al. 1998, Eberhardt 2002), with survival being the lowest and most variable for juveniles. This study did not address the causes of juvenile mortality, but it could be the result of a range of natural and human-caused hazards experienced by young and inexperienced animals. With regard to the latter, entanglement in fishing gear and vessel strike are known sources of impact (Robbins 2012, Hill et al. 2017, Stepanuk et al. 2021). Whereas adult survival was high in most years, it was slightly lower for females than males. Previous work in the GoM found lower survival of adult females in calving years and suggested a cost of the energetic burden of reproduction (Robbins 2007), but other causes, such as human interactions, warrant investigation. For example, female North Atlantic right whales with evidence of entanglement have lower survival than for males with the same level of injury (Knowlton et al. 2022). The only other area where sex-specific humpback whale survival has been studied is Gulf of St. Lawrence, Canada, and it was significantly lower there for males than females (Ramp et al. 2010). However, the population of humpback whales that summers in the Gulf of St. Lawrence differs from those in the GoM in other respects as well, such as lower overall abundance and lower female reproductive rates (Schleimer et al. 2019). Survival patterns are contextual and thus expected to differ among humpback whale populations according to the local circumstances.

The few available contemporaneous abundance estimates for this population included two minimum number alive (MNA) estimates from the same mark-recapture data used in this study and six line-transect-based abundance estimates (Table B.2, Clapham et al. 2003; Waring et al. 2007; Waring et al. 2013; Hayes et al. 2019; Palka 2020). MNA is a minimum by definition, and it depends upon the amount of sampling performed before and after each year. For this population, these MNA-based estimates were lower than the 95% probability envelope of our model-based estimates. MNA is easily calculated from mark-recapture data and avoids potentially over-estimating abundance, but may be an overly precautionary lower bound on the size of this population when information is needed in near real-time.

Comparing our results with available line-transect-based estimates is more challenging because the two methodologies estimate a different quantity: i.e., the number of animals present within a surveyed area at the time of the study versus the size of the population that uses the area. The geographic coverage is therefore an important consideration for line transect estimates, and that coverage varied somewhat from year to year. Only one such study (Palka 2020) has suggested a substantially larger abundance than estimated here (2,368, CV=0.48 in 2016). The Palka (2020) survey took place in the same season as our study, but it covered only the U.S. portion of the primary GoM feeding ground, versus also Canadian waters. That survey also included waters to the south of the GoM, and most of the sightings occurred in those more southerly waters. Although it is not possible to determine the identity of the individuals counted by Palka (2020), any GoM whales would have been included in our estimates if they had only temporarily moved out of our study area. Our surveys were not originally designed to extend south of the GoM because, historically, humpback whales were rarely observed in southerly coastal and shelf waters in summer (CETAP 1982). The Palka (2020) line-transect results add to accumulating evidence of increasing use of that area by humpback whales (Brown et al. 2018; Zoidis et al. 2021). The primary feeding ground origin of humpback whales in the New York Bight and adjacent waters is still under investigation, with only some of the individuals identified there currently known to use the adjacent waters of the GoM (Brown et al. 2022). That research will be important for determining how humpback whales along the U.S. East Coast should be managed in the future, given that these new areas potentially pose different threats.

A population growth rate of 4.6% is low for a species with a conservative average of 7.4% and a maximum plausible rate of increase of 11.8% (Zerbini et al. 2010). There are no other estimates of the population growth rate of humpback whales in the GoM or other feeding or breeding grounds in the western North Atlantic during the study period. In the 1980s, humpback whales in the GoM increased at an average annual rate of 6.5% (95%CI: 4.1-8.8, 1979–1991, Clapham et al. 2003). In the following decade, growth was either zero or 4.0%, assuming calf survival rates of 51% and 87.5%, respectively (Clapham et al. 2003). A later study found that a 51% calf survival rate was more plausible for that study period (Robbins 2007). The most recent growth rate estimate for the overall West Indies DPS was nearly a decade before this study began (1979-1993, 3.1%, Stevick et al. 2003). The results presented here suggest that population growth resumed after the 1990s at an average rate that is not distinguishable from the rate during the 1980s.

### 4.2 Insights into the recovery status of North Atlantic humpback whales

North Atlantic humpback whales were a target of historical whaling, with over 30,000 individuals killed from 1616 until the International Whaling Commission (IWC) established a moratorium for this population in 1955 (Smith & Reeves 2010). Two decades ago, the Scientific Committee of the IWC undertook a Comprehensive Assessment of the North Atlantic humpback whale to assess the level of population recovery (IWC 2001, 2002; Punt et al. 2006). Historical carrying capacity for the GoM was estimated to be 2,500-3,000 individuals, depending on model assumptions, and neither the GoM nor the North Atlantic population overall, was estimated to have reached carrying capacity (historical or current) by the year 2000 (Punt et al. 2006). However, the population dynamics model had a poor fit to some of the available data, including the limited available absolute and relative abundance trends for the GoM (Punt et al. 2006). Thus, conclusions regarding recovery status were limited and mainly served to emphasize several areas of ongoing uncertainty about population structure and other data gaps (Clapham et al. 2003; Punt et al. 2006; Bettridge et al. 2015). The present study provides new, robust results on population abundance trends in the GoM and suggests that there has been continued population growth since 2000. However, the decline in the abundance of adult females and juveniles, as well as reduced calving rates, may indicate population limitation (Eberhardt 2002). The population estimate in 2019 was at most 66% of the estimated historical carrying capacity, although our understanding of both historical and present carrying capacity for humpback whales remains limited (Punt et al. 2006). The IWC is in the process of re-evaluating population recovery in the North Atlantic (IWC 2019, 2020) and the results of the present study will help to resolve the data gaps for the GoM.

### 4.3 Monitoring a vulnerable subset of the West Indies DPS

In 2016, after 46 years as an endangered species in the United States, humpback whales that breed in the western North Atlantic, the West Indies DPS, were removed from the U.S. Endangered Species List. Assessing the status of wide-ranging oceanic populations can be challenging, and research conducted since the status change suggests that the West Indies DPS is not homogenous. The southeastern portion of the West Indies DPS exhibits different timing and preferential exchange with eastern North Atlantic feeding grounds, and documented exchange with the endangered Cape Verde Islands/Northwest Africa DPS (Stevick et al. 2016; 2018). The GoM is the feeding ground with the least exchange with the southeastern portion of the West Indies DPS (Stevick et al. 2018), but given its high level of human activity, even a small amount of exchange may ultimately prove to have management implications.

Even at the time of delisting, one of the original target goals for status assessment, population doubling over 20 years (NMFS 1991), could not be evaluated (Bettridge et al. 2015). This study focused on one portion of the West Indies DPS that has been studied on its feeding grounds for decades. The GoM feeding ground, used consistently by some members of the West Indies DPS, is an area of high human use, and the humpback whales that feed in the GoM are vulnerable to anthropogenic impacts (van der Hoop et al. 2013; Henry et al. 2022). Even for this comparatively well-studied portion of the population, impacts are likely under-estimated because they are based only on observed cases that are known, or suspected, to lead to death (e.g., Henry et al. 2022). Entanglement alone could result in an annual mortality rate of over 3% (Robbins et al. 2010). However, the present study suggests that the GoM subset of the West Indies DPS met the goal of doubling in abundance. The results further indicate that population growth remains positive, but has slowed, since the change in status under the ESA in 2016.

One of the strengths of this study is the amount and types of data available to inform the state-space model. The depth and detail of the demographic data available to this study are not common for baleen whale populations and are the product of decades of long-term research on individuals. The availability of auxiliary data to clarify the biological state of individuals that were alive but not seen is also unusual and greatly improved estimates of abundance achieved by the primary mark-recapture data alone. In the Gulf of Maine, auxiliary data come primarily from a long-standing relationship between scientists and whale-watching-based data collection programs. Those programs were historically founded by and directed by researchers, but are now largely maintained by whale-watching companies that share data with research programs. This study is an example of how formally structured collaborations of that nature can benefit science and conservation. It is challenging to study a population that ranges across an ocean basin, and population trends are still not well understood for portions of the DPS that feed in other areas of the western North Atlantic. However, the GoM is an area where population dynamics can be monitored at a high level, along with potential threats, yielding information that is particularly relevant to U.S. impacts and resource management. Such monitoring may also serve as a bellwether for DPS-wide status changes, such as from predicted climate change impacts on shared breeding habitats (von Hammerstein et al. 2022).

### 4.4 Study assumptions and limitations

This study leveraged several large, long-term data sets to estimate the abundance and trend of humpback whales in the GoM, but they were not specifically collected for this purpose. One important assumption of open population mark-recapture models is that there is no permanent emigration from the study area, which would be confounded with mortality. Humpback whale feeding grounds are largely discrete, but individuals do sometimes move among areas. Studies to date have suggested that only a small number of individuals move between the GoM and other primary feeding grounds, including individuals hypothesized to be migratory transients moving from northerly feeding grounds during shoulder seasons (Katona & Beard 1990; Stevick et al. 2006). Higher than average levels of exchange to eastern Canada in the 1980s (n=25 individuals) appeared to correlate with years of particularly low abundance of two preferred prey species in the GoM (Stevick et al. 2006). By contrast, only one such movement between these areas was detected during a subsequent ocean-basin-wide study in 1992 and 1993 (Stevick et al. 2006). In a more recent photo-identification study, 4% of individuals sighted in the GoM between 1970 and 2015 were detected on a nearby primary feeding ground (Jones 2018). No studies have formally examined whether detected movements among feeding areas are temporary or permanent. A recent study of the closest primary feeding ground (the Gulf of St. Lawrence) found declining reproductive rates during the same period as reported here (2004-2018), which the authors hypothesized to be related to resource limitations (Kershaw et al. 2021). If so, and if individual movements to other feeding grounds are resource-related, then conditions in at least the Gulf of St. Lawrence may not have been attractive for permanent emigration during that period. However, further work is required to fully understand and predict the movements of individuals between feeding grounds.

Survival rates varied across years and classes, and could conceivably include effects from permanent emigration. For example, the two periods of low apparent survival in the GoM in 2005-2006 and 2016 were proximal to documented UMEs (NOAA 2024), but the most prominent one, peaking in 2012, was not. It is possible that those deaths were not detected due to an offshore distribution of mortalities or other factors (Williams et al. 2011), as also demonstrated for North Atlantic right whales (Pace et al. 2021). However, permanent emigration is another possible explanation. Adult survival estimates were high in this study and so there is no significant concern of downward bias due to permanent movements outside of the study area. However, we cannot exclude it as possible explanation for low and variable juvenile survival and abundance. Previous ocean-scale studies of movement among primary feeding grounds in the North Atlantic are not likely to be as informative for the movement patterns of juveniles, as individuals represented only by calf-year images are not necessarily cataloged for areas outside of the GoM (Jones 2018). As noted previously, humpback whale sightings have increased immediately south of the GoM (Brown et al. 2018; Zoidis et al. 2021) where they were historically uncommon (CETAP 1982). Photo-identification studies have shown that individuals in this newly populated area are primarily juvenile whales, but not exclusively (Brown et al. 2018; Zoidis et al. 2021), including off the New York Bight apex (Brown et al. 2022). The vast majority of those individuals have subsequently returned at least once to the GoM (Center for Coastal Studies, unpublished data). Thus, while the issue of permanent emigration is a topic that warrants further investigation, particularly for low survival around 2012, at this time it does not appear to be a major source of downward bias in survival estimates, even among juveniles.

An important facet of this study was that there was an effort to detect individuals over a wide portion of the primary feeding range in the GoM. Nevertheless, all sites within the GoM were not equally sampled, and offshore sites, in particular, were challenging to reach consistently across years due to distance from shore, prevailing weather and other factors. Our results confirm previous evidence that individual humpback whales use both coastal and offshore GoM areas (Robbins 2007). Mark-recapture estimates are less vulnerable to fine-scale spatial coverage than line-transect sampling because they estimate the population that uses an area versus the number of animals present within the specific temporal and spatial bounds of a survey. Furthermore, previous research has demonstrated that there is connectivity among areas, including inshore and offshore areas, such as the southwestern GoM (Stevick et al. 2006; Robbins 2007). Biases could still potentially arise if individuals have such strong preferences for particular sites or other behaviors that result in consistent heterogeneity in capture probabilities among individuals. However, in this study, individual heterogeneity was accounted for as a parameter in the model because prior research indicated the importance of accounting for such differences, as well as sex and age, in mark-recapture studies (Hammond 1990; Robbins 2007; Ford et al. 2012; Robbins et al. 2015; Pace et al. 2017). Pace et al. (2017) confirmed through simulation that individual heterogeneity had minimal bias on true North Atlantic right whale abundance based on a similar model structure. Nevertheless, in the future, it may be useful to explore how inter-annual variation in spatial coverage for this species might affect capture probabilities.

Finally, our knowledge of the age and sex of individuals was extensive, but it was not complete. Some of the individuals that were of unknown exact age at first sighting were likely juveniles, but were treated as adults in these analyses. We took this approach because we could not allocate these individuals to a specific age class and any assignment errors would likely have resulted in an overestimate of juvenile survival and abundance. Pooling those individuals with adults could have had the effect of reducing adult survival, but as noted previously, this was more precautionary and less impactful in light of the large adult sample size. A previous study of North Atlantic right whales using a similar model and age assignment approach confirmed through simulation that the resulting adult survival estimates were close to true survival (Pace et al. 2017). However, in this study, it does still have the effect of reducing juvenile abundance, which was already low because the threshold for maturity was the earliest known age of calving. Given these facts, our results represent minimums for both juvenile abundance and the percentage of the population that was immature. These quantities are nevertheless valuable for understanding changes in demographic patterns over time.

### 4.5 Comparison to a Critically Endangered population

The Bayesian state-space mark-recapture model used here is similar to the approach recently used to estimate the abundance and trend of the critically endangered North Atlantic right whale (Pace et al. 2017). The similarity in methods invites a comparison of the dynamics of these sympatric but otherwise very different species. Right whales are a species of grave conservation concern, with a total abundance across the entire North Atlantic of only 356 individuals in 2022 and declining abundance (Linden 2023). Right whales occupy a different ecological niche than humpback whales and their temporal-spatial usage is correspondingly different. At the same time, they face the same primary types of contemporary human impacts within the GoM: entanglement and ship strikes. There are also some similarities in their population trends, despite the substantial differences in status and ecology. For example, both species generally increased in abundance until approximately 2014. After that point, right whales experienced a prolonged decline (Pace et al. 2017), whereas humpback whales declined briefly and then increased. Both species exhibited lower female survival and lower resulting female abundance (Pace et al. 2017). A notable difference between species is that juvenile humpback whales have a substantially lower average survival rate than juvenile right whales (Robbins et al. 2015; Pace et al. 2017). However, temporal trends in right whale calf survival (Reed et al. 2022) are similar to those reported here for humpback whale calves. It is conceivable that these species are facing parallel, resource-driven population limitations, although it is unlikely that the different resource trophic levels at which these two species typically feed (i.e., on copepods versus fish) would exhibit patterns on the same time scale. Both species are vulnerable to human activities, and the possibility that human impacts are simultaneously affecting the dynamics of both populations warrants more investigation.

## 5. Conclusions

Overall, our results suggest that the number of North Atlantic humpback whales in the GoM has grown in most years since 2000. The population has increased since it was removed from the Endangered Species List, but growth slowed toward the end of the study, concurrent with an on-going UME. The average population growth rate estimated here (4.6%) is moderate, both in relation to historical GoM estimates and for the species overall (Zerbini et al. 2010). Our findings of low and variable juvenile survival, declining reproductive rates and lower survival of adult females than males may indicate a population compensating for resource limitations (Eberhardt 2002). However, the population remains at no more than 66% of the pre-whaling carrying capacity estimated by Punt et al. (2006). While our understanding of current carrying capacity of the GoM remains limited, it is notable that the critically endangered North Atlantic right whale is exhibiting some of these same population trends, despite its low and declining abundance (Pace et al. 2017). Limitations on population growth can be a result of the resources available to individuals (carrying capacity), but they may also result from factors such as human impacts. One or both are possible factors affecting this population after many years of protection from commercial whaling, but with ongoing, well-established sources of human-caused mortality. The declining abundance of adult females is a potential concern here, although it is weaker than a similar effect in the critically endangered North Atlantic right whale that also uses GoM waters extensively (Pace et al. 2017). Changes in the distribution of humpback whales along the U.S. East Coast further complicate the understanding of GoM population trends, particularly for the juvenile component. Better estimates of human-caused mortality and an explicit modeling of those deaths in a population context will be important for assessing the degree to which humans are affecting the dynamics of this population.

## Acknowledgments

These data were available thanks to long-term humpback whale population research by the Center for Coastal Studies and its collaborators. The authors acknowledge the contribution of many staff, interns and volunteers at their institutions over the decades that made this work possible. They particularly thank the late C. Carlson, P. Durazo, P. Kamath, E. Kelly, A. Kennedy, T. Kirchner, B. Lynch, C. McMillan and D. Sandilands for their support and assistance. The following whale-watch companies hosted collaborating NGOs for data collection: the Dolphin Fleet Whale Watch, Captain Bill’s Whale Watch, Captain John Boats, Granite State Whale Watch, Hyannis Whale Watcher Cruises and Shearwater Excursions. Thanks also to the following for data, information or other support: Allied Whale/North Atlantic Humpback Whale Catalog, the Stellwagen Bank National Marine Sanctuary, New England Aquarium and the right whale team from the Anderson Cabot Center at the New England Aquarium. Finally, we thank the following for sharing sightings data with the GoM Humpback Whale Catalog: 7 Seas Whale Watch, Bar Harbor Whale Watch, Cape Ann Whale Watch, Coastal Research and Education Society of Long Island, Grand Manan Whale and Seabird Research Station, New England Coastal Wildlife Alliance, Newburyport Whale Watch, Quoddy Link Marine and others. The Northeast and Southeast Marine Mammal Stranding Networks provided photographs used to identify cataloged whales after death. The Atlantic Scientific Review Group’s Peer Review Panel (R. Merrick, T. McDonald, G. Nesslage, R. Kenney and J. Calambokidis) provided comments that improved the manuscript.

## Ethical statement

Primary mark-recapture surveys were undertaken under the authorization of the governments of the United States (NMFS-ESA permits 917, 755-1600, 633-1483, 633-1778, 16325 and 21485) and Canada. Auxiliary sighting data were obtained under government research permits or in accordance with local whale-watching guidelines.

## Authors’ Contributions

**Robbins**: Conceptualization, Methodology, Data curation, Formal analysis, Funding acquisition, Investigation, Project administration, Resources, Supervision, Visualization, Writing – original draft, Writing – review & editing. **Bérubé:** Data curation, Funding acquisition, Investigation, Resources, Writing – review & editing. **Clapham**: Investigation, Resources, Writing – review & editing. **Mattila:** Investigation, Resources, Writing – review & editing. **Palsbøll:** Data curation, Funding acquisition, Investigation, Resources, Writing – review & editing. **Asmutis-Silvia**: Data curation, Funding acquisition, Investigation, Resources, Writing – review & editing. **Hill:** Data curation, Funding acquisition, Investigation, Resources, Writing – review & editing. **Howes:** Data curation, Funding acquisition, Investigation, Resources, Writing – review & editing. **Landry:** Investigation, Resources, Writing – review & editing. **Lonergan:** Data curation, Funding acquisition, Investigation, Resources, Writing – review & editing. **Schulte:** Data curation, Funding acquisition, Investigation, Resources, Writing – review & editing. **Tackaberry:** Data curation, Investigation, Writing – review & editing. **Weinrich:** Data curation, Funding acquisition, Investigation, Resources, Writing – review & editing. **Pace**: Conceptualization, Methodology, Software, Formal analysis, Funding acquisition, Project management, Visualization, Writing – original draft, Writing – review & editing.

