## Supplemental Materials for "Before and after delisting: population dynamics of North Atlantic humpback whales over two decades in the Gulf of Maine"

#### Appendix A: Supplemental Methods

*Primary mark-recapture data*

The primary mark-recapture data were obtained through annual photo-identification research by the Center for Coastal Studies (CCS) in the GoM from 2000 through 2019. This work was conducted from small (10-12m) vessels operating from Nantucket, Massachusetts to Nova Scotia, Canada (Figure 1). Effort was focused on GoM areas where humpback whales tend to aggregate to feed, i.e. primarily shallow (<100-m deep) banks, ledges and slopes along the inner GoM perimeter (CETAP 1982; Payne et al. 1986; Hamazaki 2002). Between 2002 and 2004, CCS mark-recapture data were supplemented with sightings from coordinated shipboard surveys in the eastern GoM by the Northeast Fisheries Science Center (Woods Hole, MA). Humpback whale use of specific areas varies over time in response to prey availability (Payne et al. 1990; Weinrich et al. 1997). Furthermore, while individual humpback whales have site preferences they are not site-exclusive within this small feeding ground (Robbins 2007). Thus, the intent was not to sample areas *per se* but to sample sufficiently broadly across the GoM to provide a representative sample of individuals using the feeding ground in a given year. Effort was expended at each site proportional to observed densities, but not every site was surveyed nor necessarily occupied by whales in every year.

This study focused on late June through early October, a period which encompassed the peak of the feeding season and when effort was most geographically expansive. It also excluded the shoulder seasons when the population was still open to demographically staggered seasonal migration to and from its breeding grounds (Robbins 2007), and when transients from other feeding grounds were more likely to transit the GoM as part of their seasonal migration (Katona

& Beard 1991). We excluded poorly documented animals from analysis to prevent false negative matches and inflated abundance estimates (Stevick et al. 2001; Friday et al. 2008). We then compressed all primary mark-recapture based sightings of individual whales into a binary annual outcome (seen or not seen).

*Auxiliary sightings*

Small-scale data collection programs have contributed to the understanding of this population since the 1970s. In this study, they provided auxiliary data about cataloged individuals when they were not seen by the primary survey. Sightings came from a range of groups, but primarily from CCS research outside of the primary mark-recapture effort as well as from eight collaborating groups that work in localized, coastal waters off southwestern New England and in the Bay of Fundy (Figure 1). Most of these were standardized opportunistic collection programs aboard commercial whale-watching vessels operating on a near-daily basis during the seasonal data window of interest.

*Demographic data*

Demographic data for this study were obtained from the CCS Gulf of Maine Humpback Whale Catalog. The sex of an individual was determined from molecular genetic analysis of a skin sample (Palsbøll et al. 1992; Bérubé & Palsbøll 1996a, b), observation of the genital slit (Glockner 1983) and/or a calving history in the case of females. Year of birth was known for individuals cataloged as dependent calves. These individuals were categorized as juveniles until age 5, the earliest known age at first calving (Clapham 1992; Robbins 2007). Whales that were first seen after the calf year were assumed to be one year old at first sighting, but could have been older. Individuals were classified as dead only when they were matched to a carcass.

*Bayesian state-space model*

We implemented a mark-recapture statistical modeling approach similar to one employed to assess the population abundance trends of North Atlantic right whales (Pace et al. 2017; Pace 2021). We used re-sighting histories of marked individuals to estimate survival rates and abundance in a Bayesian, state-space formulation estimated using Markov Chain Monte Carlo (MCMC) simulation. Specifically, we modified the approaches of Kéry and Schaub (2011) and Royle and Dorazio (2012) to produce a multi-state formulation that relied on Jolly-Seber open population model ideas of estimating the probability of new member entry (Jolly 1965; Seber 1982) but executed it in a Bayesian framework together with data augmentation. We separated the predictive relationships associated with state transition (or biological process) from those of the observation process. The biological states modeled were: 1) not yet entered the population, 2) alive, and 3) dead. The two observational states were "seen" or "not seen". If an individual was alive prior to the start of the study, but not seen in the primary mark-recapture surveys until 2002, its observational state was coded as "not seen" in 2000 and 2001, but its biological state was coded as "alive" for those years. Individuals were also coded as alive in a given year if they were not seen in the primary survey but were known to have been alive based on auxiliary sightings. However, when the year of birth of an individual was not known, its biological state was entered as unknown (NA) until the first sighting year. An individual's state was also coded as unknown in all years after the last sighting, unless it was known to be dead.

Five juvenile age classes were defined based on the number of full years elapsed from its birth based on previously documented variation in age-specific survival rates (Robbins 2007). Individuals known to be alive for at least 5 years were considered to be adults and grouped by sex. When an individual was not first seen as a calf, it was at least one year old but could have been older. As a result, we could not confidently assign an age class until at least four years had elapsed. In this population, previous research suggests that this category of whales was predominantly juvenile when first seen (Robbins 2007), but we did not know their age with certainty. Mis-categorizing adults as juveniles would have risked inflating juvenile survival and artificially increasing population size. We therefore took a precautionary approach and placed all individuals of unknown age class into the substantially larger adult sample, recognizing that doing so might slightly depress estimates of adult survival (e.g., Pace 2021).

To estimate probability of entry into the population, which is necessary to estimate abundance, we augmented the data with sighting histories of 500 simulated individuals that could enter the population but never be seen based on estimated capture, survival and entry probabilities (Royle & Dorazio 2012).

We used logistic relationships with linear combinations of predictors (Lebreton et al. 1992) to estimate apparent survival and capture probabilities while accounting for sources of heterogeneity. Apparent survival probability was modeled for follows:

$$\mathbf{Logit(\phi_{i,t}) = \beta_1 + \beta_2*(1\text{-}sex_i)*Adult_{i,t} + \beta_3\ [Age_{i,t}] + \varepsilon_t}$$

Where: $\phi_{i,t}$ was the survival probability of the i$^{th}$ individual for the t$^{th}$ interval, $\beta_1$ was the intercept whose value in the logit is the mean of calf survival, $\beta_2$ was the added effect of being female on survival, $sex_i$ was a data value of 0 for female, 1 for male and NA for unknown, $Adult_{i,t}$ was a data value of 1 if the i$^{th}$ animal was classed as age ≥5 in the t$^{th}$ interval, $\beta_3$ was a set of factors for each age class, $Age_{i,t}$ was an index representing the age value for the i$^{th}$ individual in year t, and $\varepsilon_t$ was the random effect of year on survival. A pooled estimate of survival for the five juvenile age classes was also estimated as follows:

$$\mathbf{Logit(\phi Juv_{i,t}) = \beta_1 + \beta_2*(1\text{-}Adult_{i,t}) + \varepsilon_t}$$

Finally, we modeled capture probability based only on the primary mark-recapture data, as follows:

$$\mathbf{Logit(P_{i,t}) = \alpha_1 + \alpha_2{*}(sex_i) + Time_t + \zeta_i}$$

Where: $P_{i,t}$ was the capture probability of the $i^{th}$ individual for the $t^{th}$ year, $\alpha_1$ was the intercept and hence the effect of being an adult female on capture probability, $\alpha_2$ was the added effect of being a male on capture probability, $Time_t$ was the additive effect of the year t on average capture probability with $Time_t$=2000 was 0, and $\zeta_i$ was the random effect of the $i^{th}$ individual on capture probability.

For estimation, we assigned vague priors on all linear logistic terms except the random coefficients $\varepsilon_t$ and $\zeta_t$, as uniform(-5,5). Random coefficients $\varepsilon_t$ and $\zeta_i$ were given normal (0, δ) and normal (0, σ) priors, respectively. Standard deviation terms δ and σ were given vague priors of uniform (0.001,10). The probability of entry into the population, $\gamma_t$, was allowed to vary among time intervals, and each $\gamma_t$ was assigned a uniform (0,1) prior. Transitions among states (not yet entered, alive or dead) were modeled as a discrete categorical random variable dependent on the prior state according to the following probabilities:

| **State** | **Not entered** | **Alive** | **Dead** |
|---|---|---|---|
| Not entered | $1-\gamma_t$ | $\gamma_t$ | 0 |
| Alive | 0 | $\phi_{i,t}$ | $1-\phi_{i,t}$ |
| Dead | 0 | 0 | 1 |

The observed data (seen or not seen) were considered dependent on the animal's state and were modeled as Bernoulli(p[s]) according to the following:

| State | Seen | Not Seen |
|---|---|---|
| Not entered | 0 | 1 |
| Alive | $P_{i,t}$ | $1-P_{i,t}$ |
| Dead | 0 | 1 |

Finally, missing data on the sex of individual whales were modeled as Bernoulli(ρ), where ρ was given a somewhat informative beta (5,5). Using the above structure, data were modeled using program JAGS (Version 4.0.0) MCMC simulator (Plummer 2003) accessed via Program R version (Version 4.1, R Core Team 2012) and package "run.jags" (Version 2.0.2-8, Denwood 2016). In simulations, random starting values for each parameter were drawn from the range of its prior. Covariates concomitant with capture histories in the data augmentation set were sex=unknown (NA) and age=5+ for the adult for age class.

We provided initial values for unknown states ($state.init_{ij}$ ) which were $state.init_{ij}$= 1 prior to the first year seen and $state.init_{ij}$= 3 after the last year seen, and a value of 1 for all animals in the augmentation set of capture histories. Unknown sexes were assigned a Bernoulli (0.5) random initial value. We used an adaptation + burn in phase of 5,000 iterations and sample size of 10,000 iterations for estimation. We diagnosed convergence based on three parallel MCMC chains for which the Gelman-Rubin convergence statistic was calculated. Convergence was achieved if the convergence statistic was less than 1.1 for all model parameters (Gelman & Rubin 1992).

Annual population growth rates were calculated as $Nt+1 / Nt$ for each time step as within an MCMC chain that acted as the posterior for each growth calculation within the time series. We then used the 0.025 and 0.975 quantiles to serve as 95% credible intervals for each population growth estimate.

*Crude birth rate*

The annual crude birth rate was calculated as the percentage of calves out of all individuals seen during the primary mark-recapture surveys. Statistical evidence for a monotonic trend in the crude birth rate was based on the results of a Mann-Kendall test using Program R, package Kendall.

### Appendix B: Supplemental results

**Table B.1:** Matrix of humpback whale sightings in the primary mark-recapture survey. Each row is a sampling occasion (year). Columns indicate the total number of whales sampled in that year and the number of re-sightings in each subsequent year.

| Year | 2000 | 2001 | 2002 | 2003 | 2004 | 2005 | 2006 | 2007 | 2008 | 2009 | 2010 | 2011 | 2012 | 2013 | 2014 | 2015 | 2016 | 2017 | 2018 | 2019 |
|---|---|---|---|---|---|---|---|---|---|---|---|---|---|---|---|---|---|---|---|---|
| **2000** | 167 | 47 | 26 | 35 | 7 | 8 | 6 | 4 | 2 | 0 | 1 | 0 | 1 | 0 | 3 | 1 | 0 | 0 | 0 | 0 |
| **2001** | | 142 | 50 | 57 | 9 | 8 | 4 | 1 | 2 | 0 | 1 | 0 | 0 | 1 | 0 | 0 | 1 | 0 | 0 | 0 |
| **2002** | | | 189 | 103 | 17 | 12 | 4 | 9 | 4 | 2 | 0 | 0 | 1 | 0 | 0 | 0 | 0 | 0 | 1 | 0 |
| **2003** | | | | 404 | 119 | 111 | 38 | 24 | 20 | 5 | 4 | 0 | 3 | 2 | 3 | 0 | 0 | 0 | 0 | 1 |
| **2004** | | | | | 238 | 89 | 38 | 26 | 13 | 3 | 4 | 0 | 2 | 3 | 2 | 4 | 1 | 0 | 0 | 0 |
| **2005** | | | | | | 313 | 141 | 57 | 29 | 7 | 9 | 1 | 12 | 1 | 1 | 1 | 0 | 1 | 0 | 3 |
| **2006** | | | | | | | 289 | 135 | 60 | 20 | 7 | 3 | 6 | 8 | 6 | 5 | 1 | 1 | 1 | 2 |
| **2007** | | | | | | | | 357 | 174 | 54 | 38 | 4 | 8 | 5 | 4 | 11 | 1 | 2 | 1 | 8 |
| **2008** | | | | | | | | | 431 | 173 | 86 | 18 | 23 | 6 | 8 | 13 | 2 | 3 | 2 | 4 |
| **2009** | | | | | | | | | | 367 | 200 | 30 | 15 | 15 | 11 | 8 | 4 | 3 | 3 | 9 |
| **2010** | | | | | | | | | | | 453 | 161 | 75 | 28 | 27 | 24 | 10 | 5 | 4 | 8 |
| **2011** | | | | | | | | | | | | 255 | 145 | 23 | 10 | 12 | 4 | 1 | 2 | 2 |
| **2012** | | | | | | | | | | | | | 333 | 170 | 35 | 44 | 6 | 2 | 4 | 6 |
| **2013** | | | | | | | | | | | | | | 305 | 131 | 93 | 14 | 6 | 4 | 9 |
| **2014** | | | | | | | | | | | | | | | 298 | 189 | 15 | 8 | 12 | 11 |
| **2015** | | | | | | | | | | | | | | | | 533 | 230 | 77 | 40 | 30 |
| **2016** | | | | | | | | | | | | | | | | | 344 | 151 | 56 | 48 |
| **2017** | | | | | | | | | | | | | | | | | | 316 | 120 | 59 |
| **2018** | | | | | | | | | | | | | | | | | | | 277 | 126 |
| **2019** | | | | | | | | | | | | | | | | | | | | 390 |

**Table B.2:** Summary of previously reported abundance estimates for humpback whales in the Gulf of Maine (GoM) and adjacent waters since 1993. The GoM includes the Gulf of Maine and the Bay of Fundy.

| Year | Estimate | CV | Estimate Type | Population/Area Applicable to Estimate | Source |
|---|---|---|---|---|---|
| 1993 | 652 | 0.29 | Mark-recapture | GoM | Clapham et al. 2003 |
| 1997 | 497 | 0 | Minimum number alive | GoM | Clapham et al. 2003 |
| 1999 | 902 | 0.41 | Line-transect | GoM and Scotian Shelf[1] | Clapham et al. 2003 |
| 2002 | 521 | 0.67 | Line-transect | GoM | Palka 2006, Waring et al. 2007 |
| 2004 | 359 | 0.75 | Line-transect | GoM | Waring et al. 2007 |
| 2006 | 847 | 0.55 | Line-transect | GoM and Scotian Shelf[1] | Waring et al. 2007 |
| 2008 | 823[2] | 0 | Minimum number alive | GoM | Robbins 2010, Waring et al. 2013 |
| 2011 | 335 | 0.42 | Line-transect | GoM to Virginia | Waring et al. 2013 |
| 2015 | 896[2] | 0 | Minimum number alive | GoM | Hayes et al. 2019 |
| 2016 | 2,368 | 0.48 | Line-transect | U.S. Gulf of Maine waters to Virginia | Palka 2020, Hayes et al. 2019 |

[1]The estimate included only 25% of the Scotian Shelf abundance based on a photo-ID-based estimate of exchange with the GoM, as reported in Clapham et al. 2003.[2]Minimum number alive estimates were based on the same data set as used in this study but were calculated closer to the target year. Recalculating these values using resighting data through 2020 led to values of 915 in 2008 and 959 in 2015.

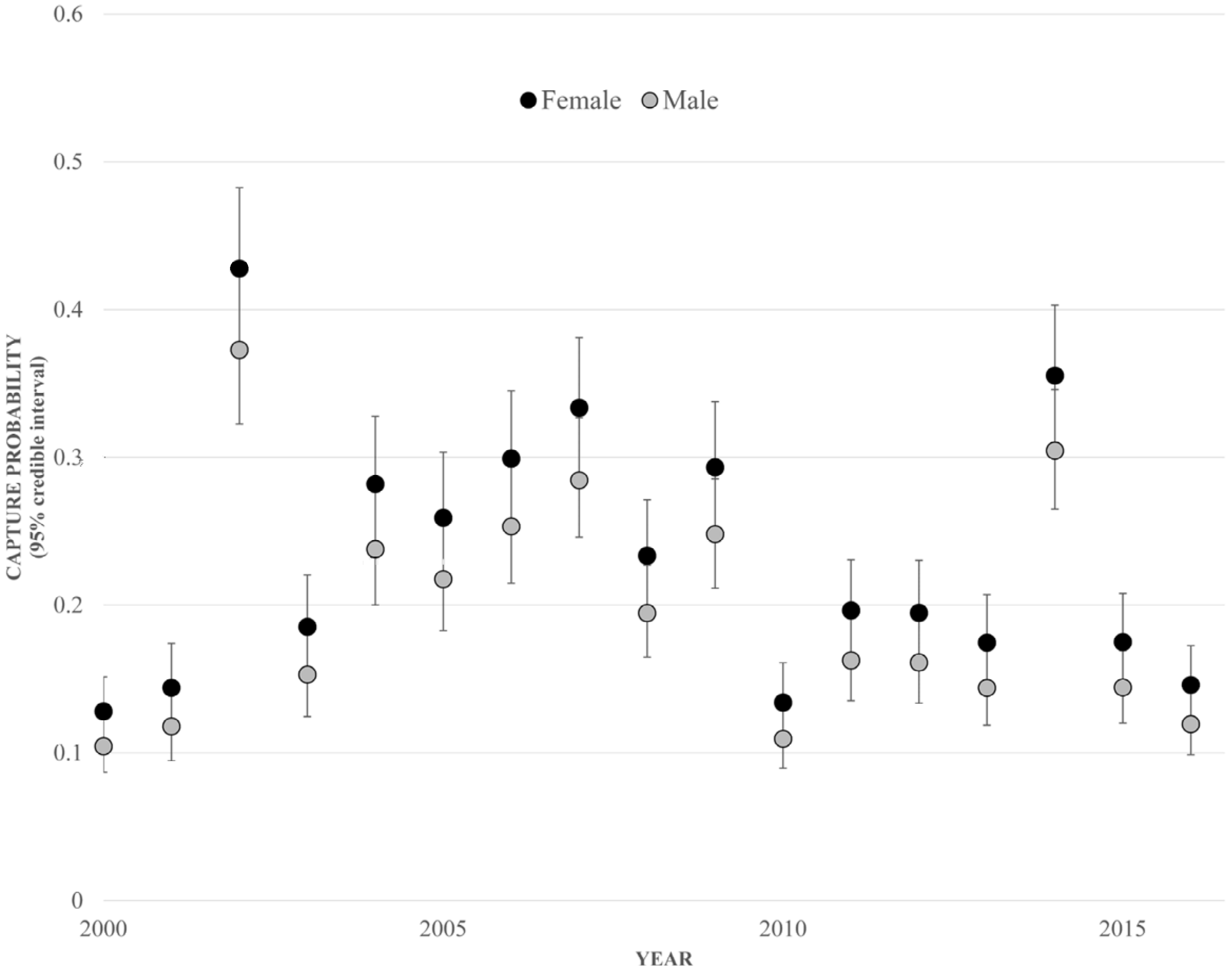


**Figure B.1:** Annual capture probabilities of humpback whale females (black circles) and males (gray circles) from 2000 through 2019 with their 95% critical regions.

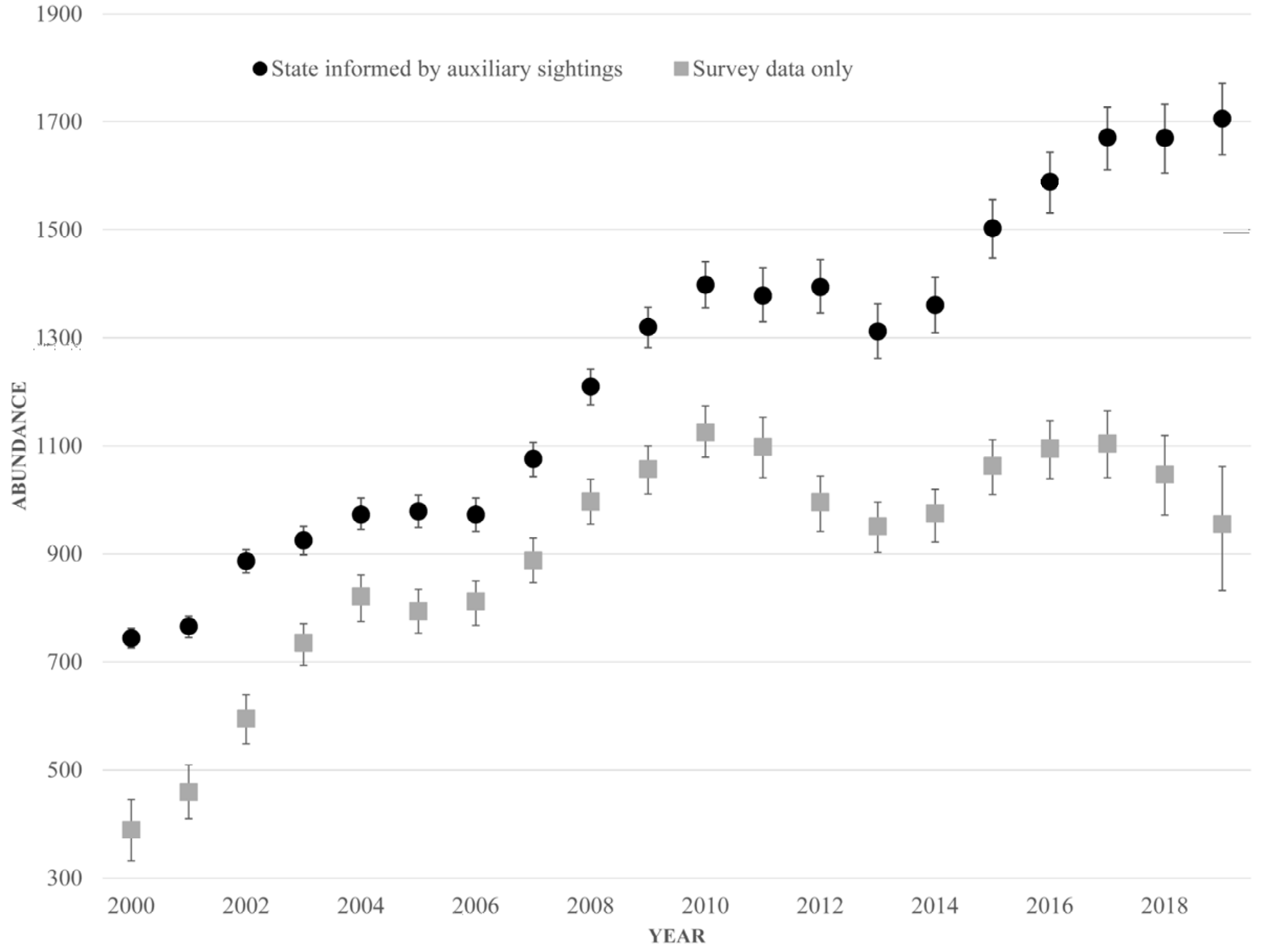


**Figure B.2:** Humpback whale abundance estimates calculated based on primary mark-recapture survey sightings alone (gray squares) and when the biological state of individuals was informed by auxiliary sightings (black circles). Error bars represent the 95% credible interval. Auxiliary data on biological state improved the estimates, particularly in the last few years of the study.
